# Reduced cutaneous CD200:CD200R1 signalling in psoriasis enhances neutrophil recruitment to skin

**DOI:** 10.1101/2022.04.01.486720

**Authors:** Holly Linley, Shafqat Jaigirdar, Karishma Mohamed, Christopher EM Griffiths, Amy Saunders

## Abstract

The skin immune system is tightly regulated to prevent inappropriate inflammation in response to harmless environmental substances. This regulation is actively maintained by mechanisms including cytokines and cell surface receptors and its loss results in inflammatory disease. In the case of psoriasis, inappropriate immune activation leads to IL-17-driven chronic inflammation, but molecular mechanisms underlying this loss of regulation are not well understood.

We reveal that immunoglobulin superfamily member CD200, and signalling via its receptor, CD200R1 are reduced in non-lesional psoriasis skin. To examine the consequences of this, CD200R1 was blocked in a mouse model of psoriasis demonstrating that the receptor limits psoriasis-like inflammation. Specifically, CD200R1 blockade enhances acanthosis, CCL20 production and neutrophil recruitment but surprisingly, macrophage function and IL-17 production were not affected, and neutrophil reactive oxygen species production was reduced.

Collectively, our data show that CD200R1 affects neutrophil function and limits inflammatory responses in healthy skin by restricting neutrophil recruitment. However, the CD200 pathway is reduced in psoriasis, resulting in a loss of immune control, and increased neutrophil recruitment in mouse models. In conclusion, we highlight a pathway that might be targeted to dampen inflammation in patients with psoriasis.

## Introduction

Barrier site immune cell activation is regulated to prevent responses against harmless environmental stimuli. This regulation is active, involving soluble mediators and cell surface receptors (1). Dysregulation of these suppressive pathways leads to chronic inflammation, such as psoriasis.

Psoriasis is a common chronic inflammatory skin disease driven by genetic and environmental factors (2). Recently, therapeutics targeting the IL-23/IL-17 axis have revolutionised the treatment of severe psoriasis (3-5) however, factors driving disease remain incompletely understood, and curative therapies are lacking. Neutrophil accumulation is a hallmark feature of psoriasis and the production of inflammatory mediators such as cytokines, ROS and hydrolytic enzymes are implicated in driving pathology (6). CCL20, is a key disease-driving chemokine, which is profoundly upregulated in psoriasis and is crucial for recruiting pathogenic T cells (7).

Psoriasis skin has inflamed lesions or plaques (PP) and non-lesional clinically normal skin (PN), which despite lacking overt inflammation, differs from healthy skin (NN). We, and others, hypothesize that PN skin is poised, ready to become inflamed if stimulated (8). We also hypothesize that this poised state is due to dysregulated immune suppressive pathways which, when intact, prevent activation of the immune system to harmless stimuli. One immune suppressive mechanism is CD200:CD200R1. CD200R1 is a cell surface receptor which detects the ligand, CD200, and activates downstream of tyrosine kinase (DOK) 1 and DOK2, leading to inhibition of MAP kinases, and thus dampening cytokine and pattern recognition receptor signalling (9, 10). CD200:CD200R1 regulates skin immunity by protecting hair follicles from autoimmune attack (11) and is required for UV-induced tolerance to contact allergens (12). At sites other than skin, CD200R1 signalling regulates responses to infectious agents (13), self-antigens (14-16) and cancers (17). Previous work showed that providing exogenous CD200 reduced psoriasis-like skin inflammation by inhibiting macrophage activity (18). However, in addition to CD200R1, murine CD200 may bind to a number of CD200R1-like receptors (19), whereas humans only possess one CD200R1-like gene which until recently was not thought to encode a functional protein (20). Therefore, investigating the role of CD200R1 in addition to CD200 is crucial for understanding the role of this receptor-ligand family in regulating the human disease, psoriasis. We hypothesize that CD200R1 signalling is dysregulated in psoriasis, allowing immune responses to occur more readily. To test this, we examined PN skin where we expect to observe changes responsible for the underlying susceptibility to psoriasis. Here we show reduced CD200, and CD200R1 signalling in PN skin which, in a mouse model of psoriasis, results in enhanced severity associated with increased skin thickening, CCL20 levels and neutrophil recruitment. Therefore, the reduced CD200 in PN skin may be an underlying factor contributing to psoriasis susceptibility.

## Results

### PN skin has reduced CD200 and CD200R1 signalling

PN skin harbours a pre-psoriatic proteomic (21) and transcriptional (22-24) profile resulting in a poised inflammatory state (8). Factors dictating this poised state are not understood, but likely involve both genetics and previous environmental insults. To determine if CD200R1 signalling is dysregulated and thus contributes to psoriasis susceptibility, CD200R1 levels were assessed in NN and PN skin by flow cytometry. PP skin was largely not examined as changes may be a consequence of inflammation rather than contributing to susceptibility. CD200R1 is expressed on most immune and non-haematopoietic (CD45-negative) cells and is similarly expressed in NN and PN skin (Figure S1). Despite similar CD200R1 levels, signalling may be dysregulated if ligand levels are perturbed. Therefore, *CD200* expression was assessed by QPCR revealing reduced expression in PN versus NN skin (Figure 1A), confirming previous RNAseq data (25). *CD200* may also be reduced in PP skin (Figure 1A), but an increased sample size is required to confirm this. By flow cytometry, CD200 was not detectable in haematopoetic skin cells (data not shown) but was seen on CD45^-^ HLA-DR^-^ and CD45^-^ HLA-DR^+^ cells (stressed keratinocytes and stem cells (26, 27)) where it was reduced in PN skin to around 50-70% of NN level (Figure 1B). Fluorescent immunohistochemistry also suggested reduced CD200 expression in PN skin, but this is not statistically significant (*p* = .12) (Figure 1C).

**Figure 1:**
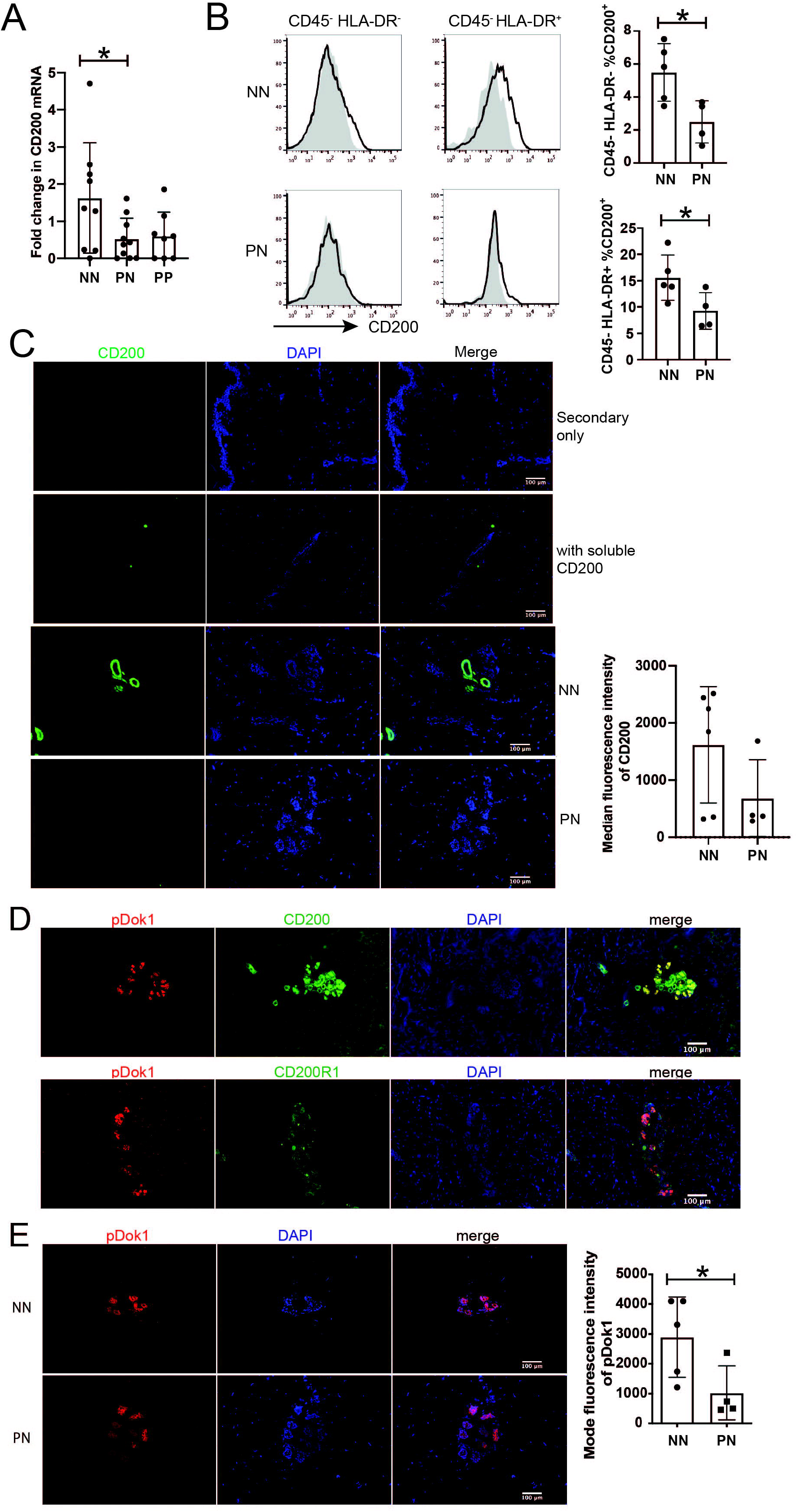
CD200 is reduced in PN skin. **A**. *CD200* QPCR on healthy (NN), non-lesional (PN) and lesional (PP) psoriasis skin relative to the mean NN value. **B**. Flow cytometry showing CD200 (black line) and isotype control (grey filled histograms) on NN and PN skin. **C-E**. Immunohistochemistry showing: **C**. CD200 in NN and PN skin with a secondary only control, and blockade of the anti-CD200 signal by prior incubation with soluble CD200, **D**. pDok1 co-staining with CD200 or CD200R1 in NN skin, **E**. pDok1 in NN and PN skin. Bar charts show all data (n = 4-6), Mean and SD shown. A was analysed by Ordinary ANOVA and Dunnett’s test. B, C and E were analysed by Mann Whitney test. * indicates *p* < .05.

CD200 staining was unexpectedly observed in interfollicular ring-like, or tubular structures. Fluorescent immunohistochemistry with markers identified these structures as eccrine sweat glands, where CD200 is expressed most highly by LGR5^+^ stem cells (Figure S2).

On engagement of CD200:CD200R1, DOK1 and DOK2 become phosphorylated leading to MAPK inhibition (9, 10, 28). To determine if the reduced CD200 in PN skin, corresponds to reduced CD200R1 signalling, pDOK was examined showing partial co-localisation of pDOK1 with both CD200 and CD200R1 (Figure 1D) and reduced pDOK1 in PN skin (Figure 1E), suggestive of reduced CD200R1 signalling.

### CD200R1 suppresses neutrophil accumulation in psoriasis-like skin inflammation

To determine consequences of reduced CD200 on skin inflammation, a mouse model of psoriasis was used, induced by topical administration of imiquimod and isosteric acid-containing Aldara cream, which induces phenotypic, histological and immunological psoriatic features (29). Mice were also intradermally injected with a CD200R1 blocking antibody (OX131) (30) on alternate days (Figure 2A). Mouse skin immune cells express CD200R1, and psoriasis-like skin inflammation largely does not affect this expression (Figure S3A). CD200R1 blockade reduced DOK1 phosphorylation (Figure 2B) as expected, which enhanced the severity of skin inflammation, shown by increased skin thickening measured using callipers (Figure 2C). Similarly, histological analysis showed Aldara cream increased epidermal thickness, which was enhanced further by CD200R1 blockade (Figure 2D-E). CD200R1 blockade also increased leukocyte numbers in draining lymph nodes (Figure 2F), and skin (Figure 2G), and most strikingly, enhanced neutrophil (Gr1^hi^ CD11b^hi^) accumulation in skin (Figure 2H-I). This CD200R1 blockade-induced increase in neutrophils is not specific to this model, as increases were also seen in an intradermal IL-23 injection model (Figure 2J).

**Figure 2:**
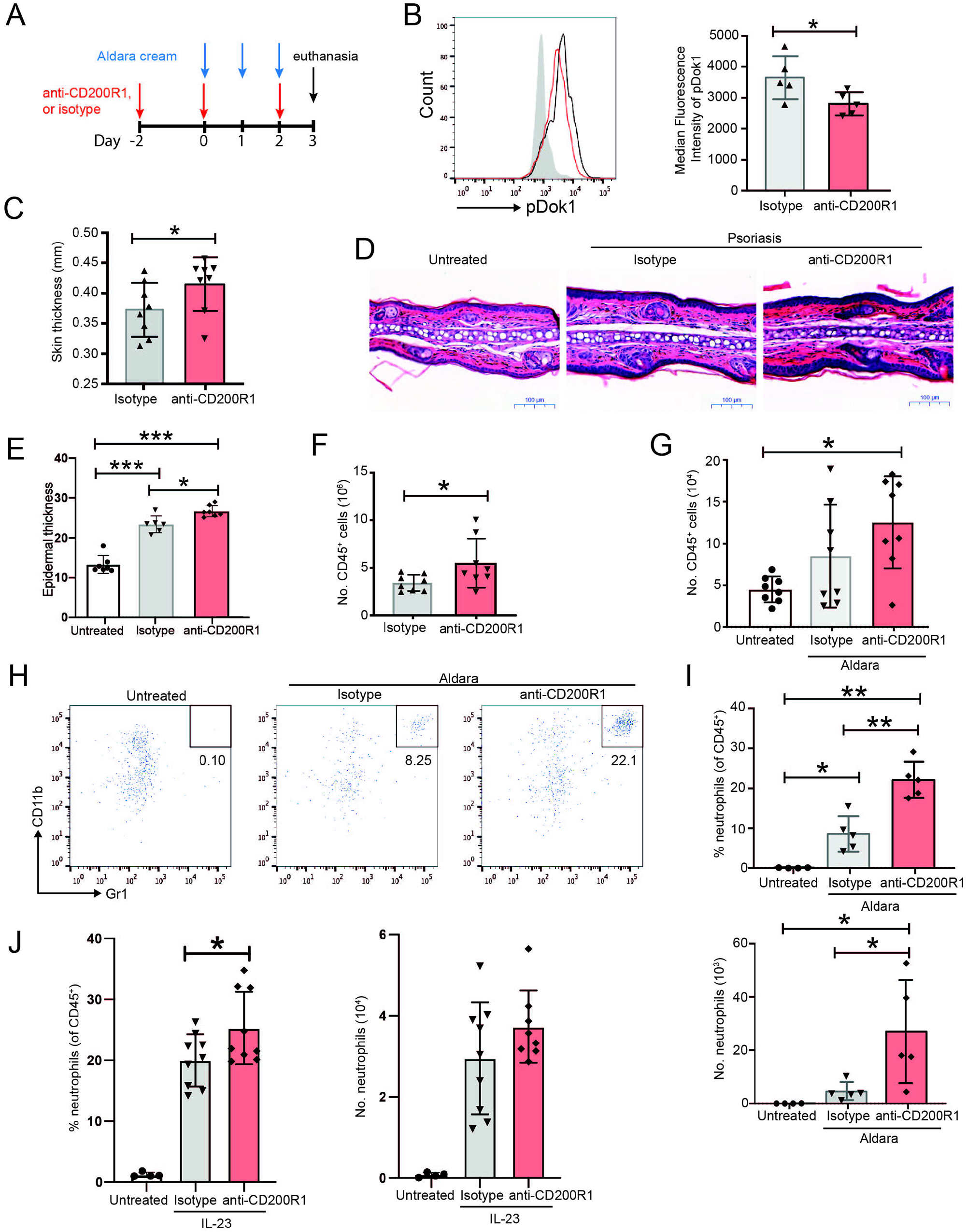
CD200R1 suppresses psoriasis-like skin inflammation by limiting neutrophil accumulation. **A-I**. Aldara cream-induced skin inflammation with CD200R1 blockade or isotype control. **A**. Timeline. **B**. pDok1 levels measured by flow cytometry in draining lymph node to avoid potential effects of enzymatic digestion which would be likely in skin. **C**. Day 3 skin thickness measured using callipers. **D**. Ear skin histology. **E**. Epidermal thickness measured on sections. **F**. Number of leukocytes in auricular lymph nodes and **G**. ear skin. **H**. Representative plots of skin neutrophils (Gr1^hi^ CD11b^hi^) within CD45^+^ cells. **I**. Proportion and number of skin neutrophils. **J**. Proportion and number of neutrophils in skin in the intradermal IL-23 injection model. B, H, I, data from one representative experiment (n = 5 for each independent experiment). C-G, J, data pooled from two independent experiments (n = 6-10). Mean and SD shown. Analysed using unpaired t tests (for 2 groups of data, B, C, F) or Brown-Forsythe and Welch ANOVA (for >2 groups of data, E, G, I, J) with Dunnett’s test. * indicates *p* < .05, ** indicates *p* < .01, *** indicates *p* < .001.

### CD200R1 blockade does not enhance imiquimod-induced macrophage activity

CD200:CD200R1 signalling is known to suppress macrophage cytokine production (31-35). Therefore, the effect of CD200R1 blockade on macrophages was measured in this model. Macrophage numbers, and IL-23 and TNFα production were not affected by skin inflammation, whereas co-stimulatory molecule expression and macrophage IL-1β and IL-6 production were induced (Figure 3A-B). Unexpectedly, CD200R1 blockade did not affect any of these parameters suggesting that CD200R1 does not affect macrophage function in this model (Figure 3A-B).

**Figure 3:**
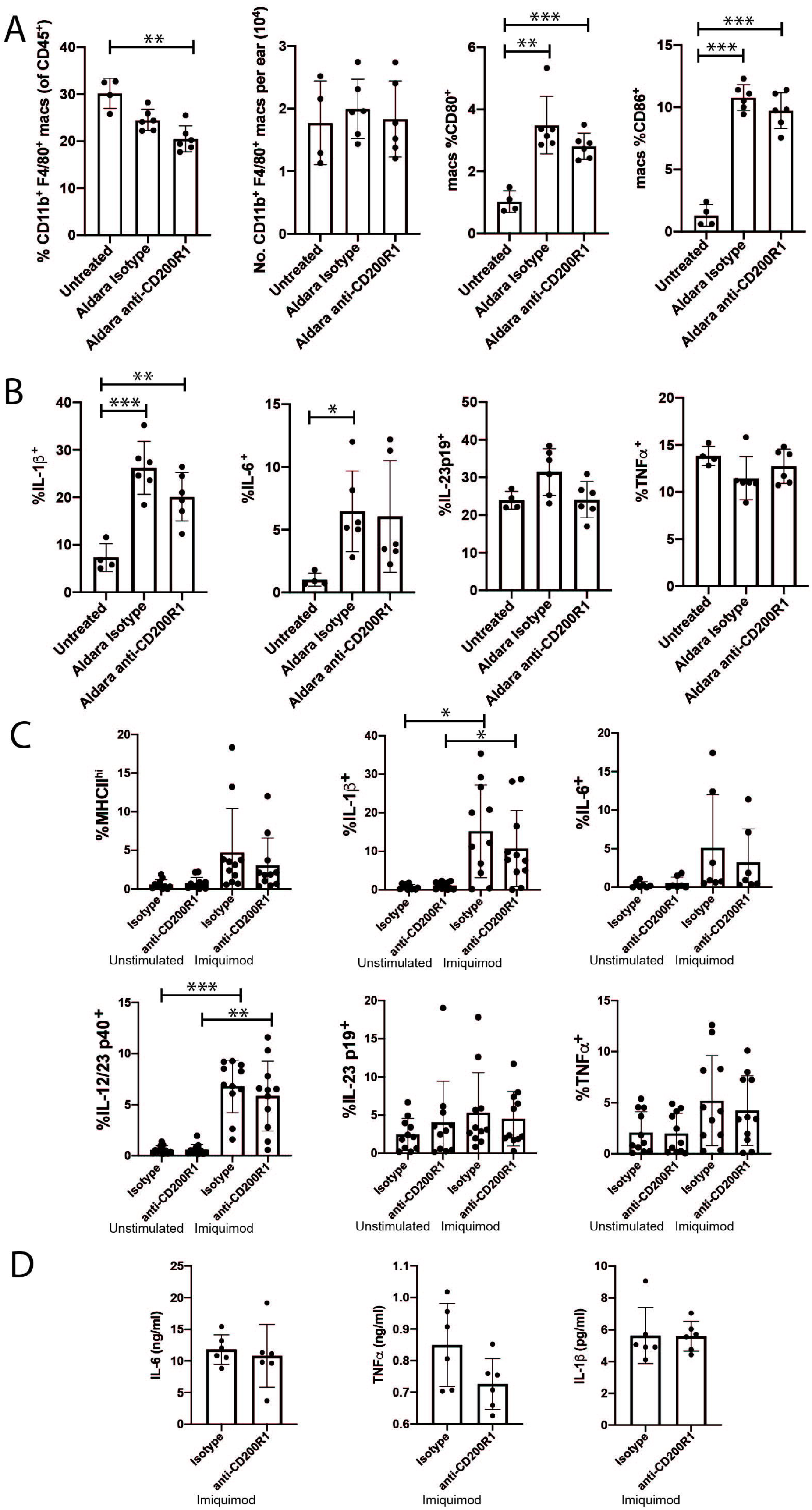
Blocking CD200R1 does not affect macrophage responses to Aldara cream or Imiquimod. **A-B** Aldara cream-induced psoriasis-like skin inflammation with CD200R1 blockade or isotype control. **A**. Skin macrophage numbers and costimulatory molecule expression. **B**. Proportion of skin macrophages producing cytokines, measured by flow cytometry. **C-D**. Imiquimod stimulated BMDM with CD200R1 blockade, or isotype control. Cytokine production by flow cytometry **C**., and ELISA, **D**. Each data point is from a separate (individual mouse) BMDM culture. Data pooled from 2-3 independent experiments (n = 4-11). Mean and SD shown. Analysed using unpaired t tests (for 2 groups of data, D) or Brown-Forsythe and Welch ANOVA (for >2 groups of data, A-C) with Dunnett’s test. * indicates *p* < .05, ** indicates *p* < .01, *** indicates *p* < .001.

Previously, CD200R1 agonists were shown to reduce IFNγ or IL-17-stimulated pro-inflammatory cytokine production by peritoneal macrophages, but no effect was seen on lipopolysaccharide (LPS)-stimulated cells (31). To determine how CD200R1 affects imiquimod-stimulated cytokine production, bone marrow-derived macrophages (BMDM) were stimulated with imiquimod in the presence of CD200R1 blockade. Imiquimod significantly induced IL-1β and IL-12/23p40 but CD200R1 blockade did not affect this, MHC class II expression, or IL-6, IL-23p19 or TNFα production (Figure 3C-D), demonstrating again no effect of CD200R1 blockade on imiquimod-induced macrophage activity.

### CD200R1 blockade does not affect IL-17 production by innate lymphoid cells (ILCs) or dermal γδ T cells

To determine mechanisms by which CD200R1 blockade affects psoriasis-like skin inflammation, cytokine levels were measured in inflamed ear tissue. Surprisingly, CD200R1 blockade, did not significantly alter cytokine levels (Figure 4A). Psoriasis, and the mouse models used here are driven by IL-23/17 with IL-17 primarily produced by γδ T cells with a contribution from ILCs (36). CD200R1 blockade had no effect on IL-17 production by either cell type, however IL-17 was also not significantly induced in either psoriasis model (Figure 4B-C) at these time points. CD200R1 is expressed by the majority of ILCs and a small proportion of CD3^low^ γδ T cells in skin (Figure 4D-E), suggesting a possible direct role. Therefore, to determine if CD200R1 blockade effects IL-17 production, *in vitro* IL-23 stimulations were performed. CD3^low^ γδ T cells and ILCs produced IL-17 in response to IL-23, but CD200R1 blockade had no effect on this (Figure 4F-G), confirming that CD200R1 blockade promotes psoriasis-like skin inflammation in an IL-17-independent manner.

**Figure 4:**
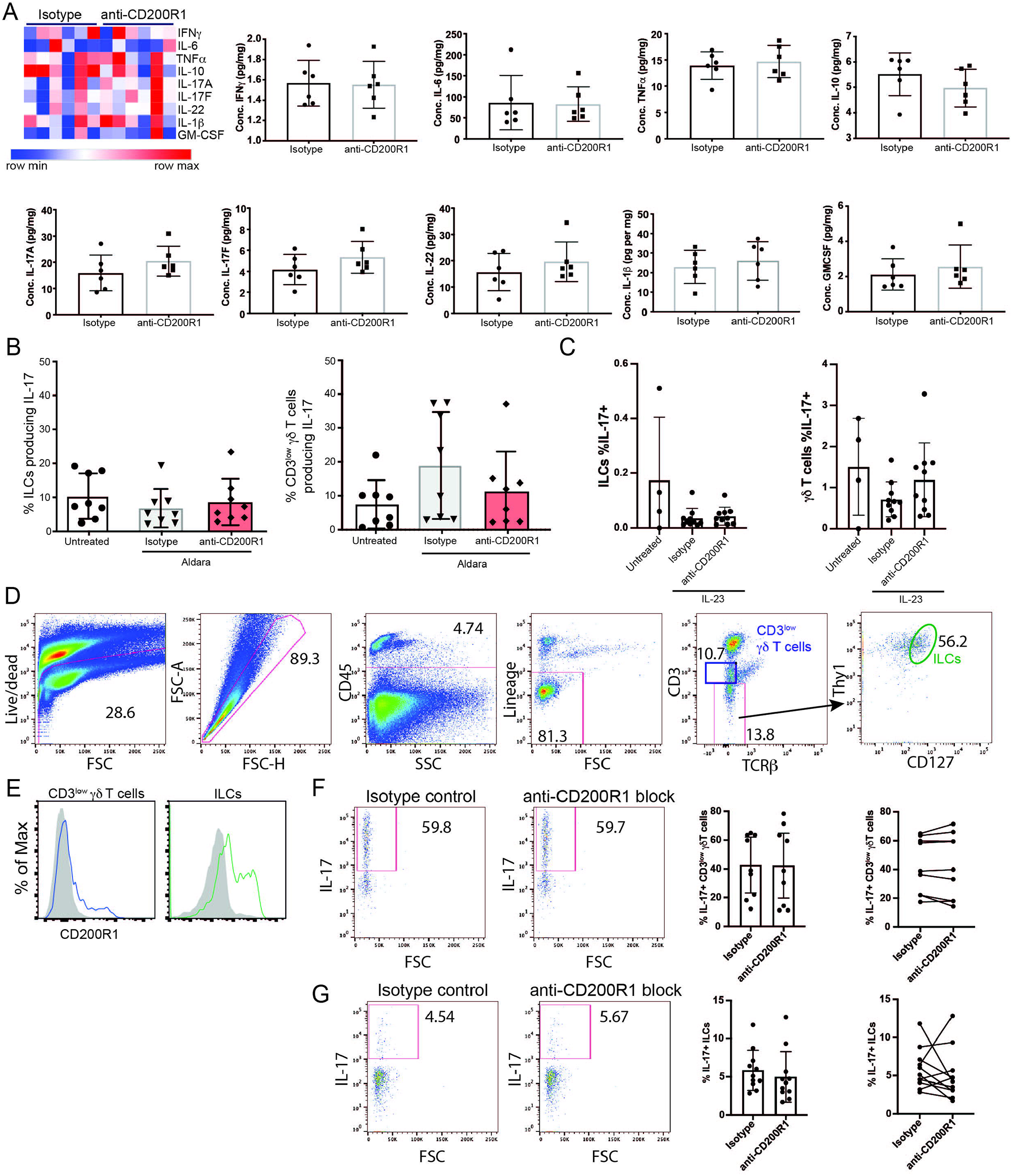
Blocking CD200R1 does not affect IL-17 production. **A-B** Aldara cream-induced skin inflammation with CD200R1 blockade or isotype control. **A**. Cytometric bead array measuring skin cytokines. **B**. IL-17 production in draining lymph node cells measured by flow cytometry. **C**. IL-17 production in lymph node of intradermal IL-23 injection model. **D**. Gating strategy for skin CD3^low^ γδ T cells (blue) and ILCs (green). **E**. CD200R1 expression in mouse skin cells. **F**. IL-17 production by IL-23 stimulated mouse skin CD3^low^ γδ T cells and, **G**. ILCs. Data pooled from 2-3 independent experiments, (n = 6-11). Mean and SD shown. Analysed using unpaired t tests (for 2 groups of data, A, F) or Brown-Forsythe and Welch ANOVA (for >2 groups of data, B, C) with Dunnett’s multiple comparison test. *p* > .05.

### CD200R1 blockade enhances neutrophil recruitment but inhibits ROS production

As CD200R1 blockade increases neutrophil accumulation in inflamed skin (Figure 2H-J), the effect of CD200R1 on neutrophils was examined. CD200R1 is expressed on inflamed skin neutrophils (Figure 5A), suggesting CD200R1 blockade may directly affect neutrophils. Neutrophils are rare in uninflamed skin, so these cells were not examined here. Neutrophils are considered terminally differentiated and traffic to sites of inflammation to carry out effector functions before undergoing cell death. Therefore, to determine if CD200R1 blockade increases neutrophils by decreasing cell death, AnnexinV/7AAD staining was used. CD200R1 blockade had no effect on neutrophil apoptosis or cell death (Figure 5B), suggesting CD200R1 blockade may instead increase neutrophil recruitment. To examine this, CXCL1 was administered intradermally which enhanced neutrophil recruitment by around 5-fold over the PBS control. CD200R1 blockade further enhanced this (Figure 5C) demonstrating that CD200R1 restricts neutrophil recruitment to skin.

**Figure 5:**
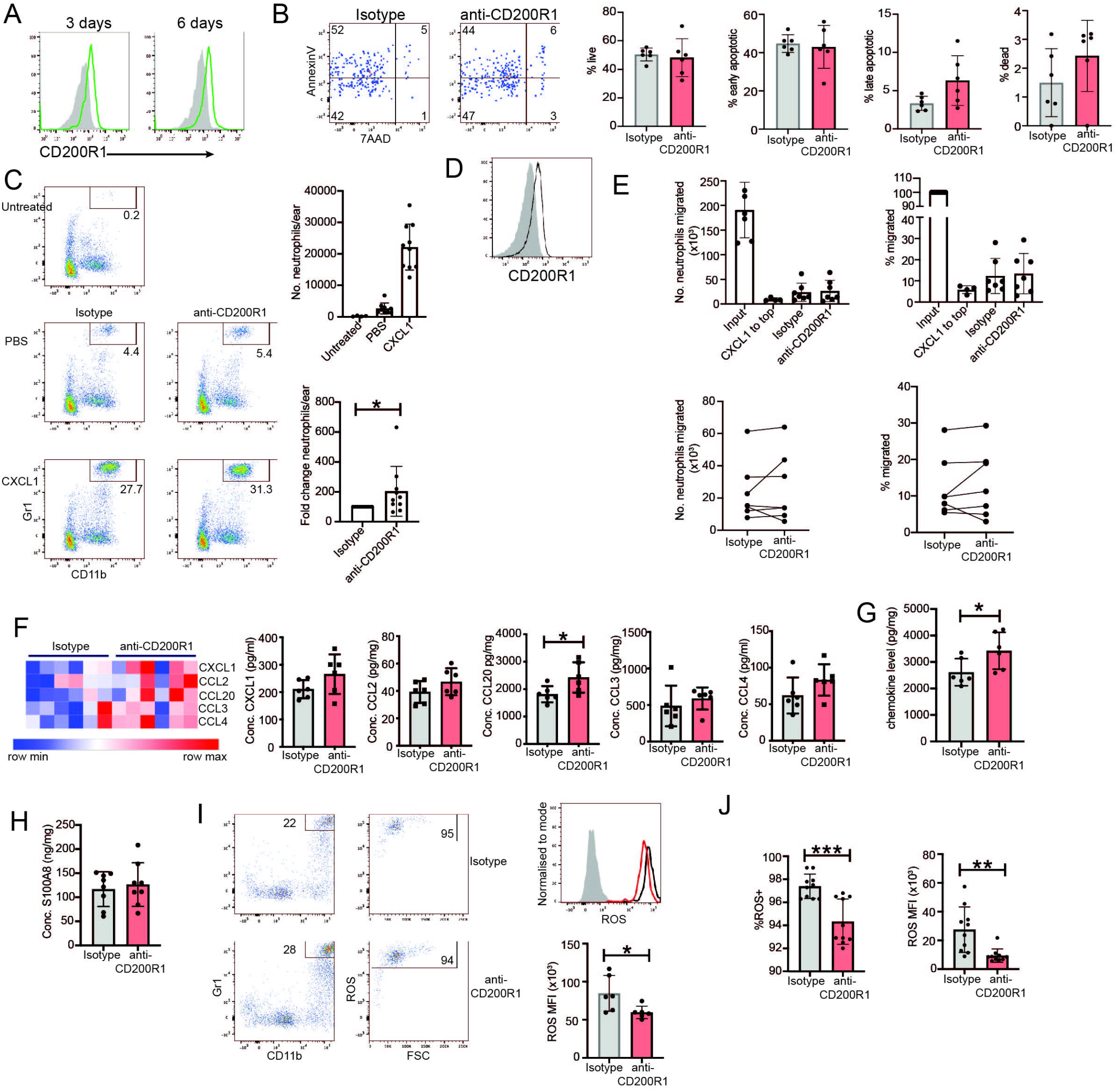
CD200R1 blockade promotes neutrophil recruitment into the skin and is associated with increased CCL20, but reduced ROS production. **A**. CD200R1 expression on inflamed skin neutrophils from skin treated with Aldara cream for the indicated days. **B-C, F-I**. Aldara cream-induced skin inflammation with CD200R1 blockade or isotype control. Neutrophil: **B**. apoptosis and cell death, **C**. *in vivo* recruitment to intradermal CXCL1 administration. **D**. CD200R1 expression on bone marrow neutrophils. **E**. BM neutrophil migration towards CXCL1 in transwell assays. ‘Input’ is cells recovered from a well lacking a transwell chamber. CXCL1 was added to the top chamber as a negative migration control, or to the bottom chamber to stimulate migration in the presence of the isotype control, or anti-CD200R1. **F-G** Skin chemokines measured by cytometric bead array. **H**. S100A8 ELISA on skin extracts. **I**. Neutrophil ROS production. Filled histogram-fluorescence minus one control, black line-isotype, red line-CD200R1 blockade. **J**. Neutrophil ROS in the intradermal IL-23 injection model. MFI: median fluorescence intensity. Data are pooled from 2-4 independent experiments (n = 6-10). Mean and SD shown. Analysed using unpaired t test (for 2 groups of data, B-D, F-J) or Brown-Forsythe and Welch ANOVA (for >2 groups of data, E) with Dunnett’s multiple comparison test. * indicates *p* < .05, ** indicates *p* < .01, *** indicates *p* < .001.

Similar to skin, bone marrow neutrophils express CD200R1 (Figure 5D) and the effect of CD200R1 blockade on transwell migration was measured using these cells due to ease of isolation. CXCL1 induced migration, but CD200R1 blockade did not affect this (Figure 5E), suggesting that CD200R1 does not directly affect neutrophil migration outside of tissues. Therefore, either CD200R1 blockade directly promotes neutrophil migration but only in tissues, or this is an indirect effect.

Chemokines direct cell recruitment, therefore, the effect of CD200R1 blockade on chemokine levels were measured and showed no significant changes in CXCL1, CCL2, CCL3 or CCL4 levels, but an increase in CCL20 and cytokines overall (Figure 5F-G). S100A8, a chemoattractant peptide particularly potent at recruiting neutrophils, and associated with psoriasis (37) was not affected by CD200R1 blockade (Figure 5H). Therefore, CD200R1 blockade increases CCL20, which is likely responsible for increased neutrophil recruitment similar to the reported recruitment of neutrophils to the central nervous system and *in vitro* (38).

A key neutrophil function is ROS production, which plays a role in the pathogenesis of psoriasis and is crucial for killing extracellular pathogens (39). Surprisingly, CD200R1 blockade reduced neutrophil ROS production in both psoriasis models (Figure 5I-J), in accordance with previous work (40). Together these data show that CD200R1 blockade modestly enhances skin thickening and vastly increases neutrophil recruitment in psoriasis models. Therefore, the reduced CD200 and CD200R1 signalling in PN skin will likely lead to increased neutrophil accumulation in response to a challenge, enhancing the immune response and potentially promoting the onset of a psoriasis flare.

## Discussion

Hair follicle stem cells express CD200 (41) which enforces immune privilege (11). Here we demonstrate previously unknown expression of CD200 in eccrine sweat gland stem cells. As sweat glands are a microbial niche (42), and their stem cells contribute to repair (43, 44) CD200 may also enforce immune privilege here, although this remains to be examined.

We show reduced CD200 in PN skin, similar to previous RNAseq data (25). Conversely, recent studies showed elevated soluble CD200 in psoriasis patient blood (45, 46), tempting speculation that the reduced cell-associated CD200 in skin, may be caused by enhanced CD200 cleavage. However, the reduced *CD200* mRNA levels seen (Figure 1A and previously (25)), suggest that reduced CD200 in skin is due (at least in part) to decreased transcription.

CD200R1 blockade is shown here to increase neutrophil recruitment (Figures 2 and 5), which is potentially caused by elevated CCL20 levels (Figure 5F). The receptor for CCL20, CCR6, is expressed on activated neutrophils (47), and CCL20 can directly attract neutrophils (38), however, it remains unknown if this is a direct effect of CCL20 on neutrophils, which cells produce CCL20 and how CD200R1 blockade increases CCL20 levels. Previous work showed CD200R1KO mice have increased lung neutrophils in *F. tularensis* infection (40) suggesting that suppression of neutrophil recruitment is a CD200R1 function across multiple barrier tissues and inflammatory conditions.

Exogenous CD200 dampens inflammation in a similar murine psoriasis model (18), suggesting there may be therapeutic benefit to manipulating this pathway. However, in contrast to our data, systemic CD200 provision reduced cytokines in a psoriasis model and cultured macrophages (18). In contrast, here CD200R1 blockade had no effect on macrophage activity (Figure 3), suggesting CD200R1 blockade and exogenous CD200 do not give opposite outcomes. This may reflect differences in blocking versus activating this pathway, or it may suggest that exogenous CD200 engages the CD200R-like receptors in addition to CD200R1.

Although the CD200:CD200R1 pathway is associated with reduced macrophage pro-inflammatory cytokine production, data demonstrating this mainly use CD200R1 agonists (31-35), and there is a lack of data from CD200R1-deficient macrophages or macrophages with inhibited CD200:CD200R1. Indeed, one report showed reduced cytokine production from CD200R1KO macrophages (48). Therefore, it is unclear if CD200R1 and CD200 affect macrophage cytokine production similarly. Certainly, blocking CD200R1 did not affect imiquimod-induced cytokine production (Figure 3), but it remains unclear if this is imiquimod-specific, or if CD200 has CD200R1-independent affects. Recently, CD200R1 was shown to potentiate TLR7/8-induced IFNγ production if cells were pre-treated with IFNα (49), suggesting the environmental milieu may influence CD200R1 function, adding further complexity.

Our data here demonstrate that immune suppressive pathways are dysregulated in PN skin. Specifically, CD200:CD200R1 signalling is reduced which promotes the recruitment of neutrophils in mouse models. Therefore, targeting CD200R1 signalling may be a novel therapeutic strategy for treating psoriasis, which warrants further investigation. Given the success of blocking immune suppressive pathways in other areas of medicine, for example checkpoint inhibitors in cancer, this approach may be highly advantageous for psoriasis treatment.

## Methods

### Human tissue

Experiments were performed in accordance with the Declaration of Helsinki and informed consent was obtained. 6 mm punch biopsies of photo-protected buttock skin tissue were taken from NN, PP or PN (>5 cm from a lesion) skin (ethics NW10/H1005/77). Volunteers (demographic data in Supporting Information Table S1) were excluded for systemic immunosuppressive, or topical medication use within two weeks of biopsy. Abdominoplasty skin for immunohistochemistry and optimizing flow cytometry was obtained through the Manchester Skin Health Biobank (ethics NW09/H1010/10).

### Human skin cell isolation

Skin was digested with 1 mg/ml Dispase II (Roche) to split epidermis and dermis before digestion with 0.5 Wunch units/ml Collagenase (Roche, Liberase TM) for 6 hr (epidermis) or 18 hr (dermis).

### Flow cytometric analysis of human skin cells

Skin cells were incubated with 50 µg/ml human IgG (Sigma), Near IR Dead cell stain (Invitrogen) and fluorescent antibodies (Supporting Information Table S2) before fixation with Foxp3/Transcription Factor Buffer Staining Set (eBioscience), and analysis using a BD Fortessa flow cytometer and FlowJo (TreeStar).

### Fluorescent Immunohistochemistry

Fresh frozen skin sections (7 µm) were fixed in cold acetone, permeabilized with Triton X-100 and blocked with 1% BSA in TBS prior to antibody incubations (Supporting Information Table S3). Where Tyramide signal amplification reagent (Invitrogen) was used, endogenous peroxidase activity was pre-quenched with 2% H_2_O_2_. Co-stained sections were stained sequentially and were quenched with 15% H_2_O_2_ before the second antibody stain. On occasion, anti-human CD200 was pre-incubated with a 1.5-fold molar amount of human CD200Fc (R&D Systems) prior to staining, to check specificity. Prolong Diamond antifade reagent with DAPI (Invitrogen) was used for mounting. Images were acquired using a Zeiss Axioimager.D2 microscope and captured using a Coolsnap HQ2 camera (Photometrics) using MetaVue Software (Molecular Devices). Images were processed and analyzed using ImageJ. Fluorescence intensity was measured using ImageJ and tracing the regions of interest where measurements were taken.

### RNA extractions and QPCR

RNA was extracted from ≥ 4 skin sections per sample using Pure Link RNA Mini Kit (fresh frozen sections), or Pure Link FFPE RNA Isolation Kit (ThermoFisher Scientific) (fixed paraffin embedded sections). cDNA was synthesized using High capacity RNA to cDNA kit (Invitrogen), and qPCR was performed using Fast Sybr Green Master Mix, and QuantStudio 12k flex real time PCR system (Invitrogen). Primers used: GAPDH For: ATCAGCAATGCCTCCTGCAC, GAPDH Rev: TGGCATGGACTGTGGTCATG, hCD200 For: CCTGGAGGATGAAGGGTGTTAC, hCD200 Rev: AGTGAAGGGATACTATGGGCTGT. Primers designed by Origene and span at least one intron-exon boundary. Data were analysed by 2^-ΔΔCT^ method, comparing each sample to the average value of the healthy samples.

### Mouse skin inflammation models

All animal experiments were ethically approved and performed in accordance with the UK Home Office Animals (Scientific Procedures) Act 1986 under project license P925B5966. Male C57BL/6 mice (Charles River) were used at 7 to 10 weeks. Ears were treated topically with 20 mg Aldara cream (Meda Pharmeceuticals, which contains Imiquimod and isosteric acid (50)) daily, for 3 days and skin thickness was measured by digital micrometer (Mitutoyo). Two days prior to Aldara cream treatment, and on days 0 and 2 of Aldara cream treatment, intradermal injections of 2.5 µg anti-CD200R1 (OX131, Absolute Antibody), or rat IgG_1_ isotype control (eBioscience) were given. On day 3 mice were euthanized and ear skin and draining lymph nodes (auricular) were analysed. This short time course was used as C57BL/6 mice become relatively highly inflamed, therefore longer treatment regimens were not necessary for the measures of inflammation used here.

For the IL-23-induced model, 1 µg recombinant mouse IL-23 (Biolegend) was intradermally injected per ear, daily for 5 days. Two days prior to IL-23 treatment, and on days 0, 2 and 4 of IL-23 treatment, the mice were intradermally injected with anti-CD200R1 or isotype control as described above. On day 5 mice were euthanized and ear skin and draining lymph nodes (auricular) were analysed.

### Mouse cell isolation

Ears were split and digested with 0.8% w/v Trypsin (Sigma) for 30 min, then chopped and digested in 0.1 mg/ml (0.5 Wunch units/ml) Liberase TM (Roche) at 37 °C for 1 hr. Dorsal skin was similarly digested, but with 1 mg/ml Dispase II (Roche) instead of Liberase. Skin and auricular lymph node cells were passed through 70 µm cell strainers and counted.

### Bone Marrow Derived Macrophages

Bone marrow cells were isolated from femur and tibia, red blood cells were lysed with ACK lysis buffer (Lonza) and cells were plated at 5 × 10^5^ cells/10 ml with 20 ng/ml M-CSF (PeproTech). On day 6 BMDM were replated at 1 × 10^6^ cells per well in 24-well plates and stimulated with 10 µg/ml Imiquimod (Biolegend) with 10 µg/ml anti-CD200R1 (OX131, Absolute Antibody), or rat IgG_1_ isotype control (eBioscience) for 20 hrs. Purity was assessed by flow cytometry and cells were typically 90% F4/80^+^ CD11b^+^ (Figure S3B).

### Flow cytometric analysis of mouse cells

Cells were incubated with 0.5 µg/ml anti-CD16/32 (2.4G2, BD Bioscience), Near IR Dead cell stain (Invitrogen) and fluorescently labelled antibodies (see Supporting Information Table S4). For cytokine analysis, cells were cultured for 4 hr with 10 µM Brefeldin A prior to staining. Cells were fixed and permeabilized with Foxp3/Transcription Factor Buffer Staining Set (eBioscience). For apoptosis the Annexin V Apoptosis Detection Kit eFluor 450 (Invitrogen) was used. ROS was assayed using Total Reactive Oxygen Species (ROS) Assay Kit 520 nm (Invitrogen). For pDok cells were fixed with BD PhosFlow Lyse/Fix buffer then Foxp3/Transcription Factor Buffer Staining Set (eBioscience) before a permeabilization with BD PhosFlow Perm Buffer III and staining with pDok1 Y398, then anti-rabbit AF488. Lymph node cells were stained rather than skin to avoid potential effects of lengthy enzymatic digestion of skin on pDok1 levels.

Cells were analysed using a BD Fortessa or LSRII flow cytometer and FlowJo (TreeStar).

### Cytokine and chemokine quantification

Skin was chopped, frozen at -80 °C, pulverized with a Tissuelyser II (QIAGEN) and lysed with 1% Triton X-100 (Sigma) with cOmplete Mini protease inhibitors (Roche). Total protein was quantified by Pierce BCA Protein Assay kit (Thermo Scientific). Analytes were quantified using LegendPlex mouse Th17 Panel (IFNγ, TNFα, IL-6, IL-10, IL-17A, IL-17F, IL-21, IL-22) and mouse Pro-inflammatory Chemokine Panel (CCL20, CXCL1, CCL2, CCL3, CCL4) (Biolegend) using a FACSVerse flow cytometer (BD Bioscience). S100A8 was measured by ELISA (Bio Techne).

### H&E staining

Skin was fixed in 10% neutral buffered formalin, embedded in paraffin and cut to 5 µm. Haematoxylin and eosin staining was performed by Shandon Varistain V24-4. Image acquisition used 3D-Histech Pannoramic-250 microscope slide-scanner and Case Viewer software (3D-Histech).

### *In vitro* γδ T cell and ILC activation assay

Mouse dorsal skin cells were stimulated with 40 ng/ml recombinant IL-23 (Biolegend) for 20 hr, with 10 µg/ml anti CD200R1 (OX131, Absolute Antibody) or isotype control (rat IgG_1_, eBioscience).

### CXCL1 intradermal neutrophil recruitment assay

Ears were intradermally injected with 2.5 µg anti-CD200R1 (OX131, Absolute Antibody) or isotype control (rat IgG_1_, eBioscience) in PBS or 1 µg CXCL1. After 3 hr ear cells were analysed by flow cytometry for neutrophils.

### Neutrophil Transwell migration assay

Neutrophils were isolated from bone marrow using Histopaque gradient centrifugation (51), placed in 3 µm transwell inserts with 10 µg/ml anti-CD200R1 (OX131, Absolute Antibody), or rat IgG_1_ isotype control (eBioscience) and 20 nM CXCL1 was added to the bottom chamber. After 1hr cells in the bottom chamber (migrated) were analysed by flow cytometry with precision count beads (Biolegend).

### Statistics

Graphs were plotted using GraphPad prism. Normality was tested by Shapiro Wilk test. Mann Whitney tests were used for human data with 2 groups. Ordinary ANOVA with Dunnett’s multiple comparison test was used for human data with more than 2 groups (as SD were equal). Mouse data were all normally distributed so unpaired t tests (for 2 groups of data) or Brown-Forsythe and Welch ANOVA (for >2 groups of data) with Dunnett’s multiple comparison test were used. Data points show data from one biological sample (1 individual).

## Abbreviations

BMDM: Bone marrow-derived macrophages
DOK: Downstream of tyrosine kinase
ILC: innate lymphoid cell
NN: healthy
PN: non-lesional psoriasis
PP: psoriasis plaque
ROS: reactive oxygen species

## Data Availability Statement

The data supporting the findings of this study are available from the corresponding author upon reasonable request.

## Author contributions

HL, SJ and KM performed experiments, CEMG assisted with experimental design and clinical sample procurement, AS conceived project, performed experiments and wrote manuscript.

## Acknowledgments

Research was funded by a pre-competitive, open innovation award to the Manchester Collaborative Centre for Inflammation Research, by University of Manchester, AstraZeneca and GSK, and a Wellcome Trust and Royal Society, Sir Henry Dale Fellowship to AS (109375/Z/15/Z). CEMG is part funded by the NIHR Biomedical Research Centre.

The University of Manchester Bioimaging Facility microscopes used in this study were purchased with grants from BBSRC, Wellcome and the University of Manchester Strategic Fund.

We acknowledge assistance from Peter Walker, Roger Meadows and Gareth Howell, and the use of the University of Manchester Histology, Flow Cytometry, and Biological Services facilities. Human skin biopsies were obtained with assistance from Jean Bastrilles, Gill Aarons and Marie Durkin, abdominoplasty samples were obtained through the Manchester Skin Health Biobank. Skin biopsy sections for QPCR were provided by Rajia Bahri and Silvia Bulfone Paus.

