## Supplementary Information for "Reduced cutaneous CD200:CD200R1 signalling in psoriasis enhances neutrophil recruitment to skin"

**Supporting Information**

**CD200 is expressed in the eccrine sweat gland**

CD200 is expressed in stem-like cells in the hair follicle (1, 2), but expression in intra-follicular regions has not been well characterised. The buttock skin used to examine CD200 and CD200R1 levels is relatively devoid of hair follicles, yet CD200 and CD200R1 expression was observed in tubular structures (Figures 1 and S1). To determine the location of this intrafollicular expression, fluorescent immunohistochemistry was used. Co-staining for CD200 and the vascular endothelial marker, CD31, and the lymphatic marker, LYVE-1, showed some co-staining, but also distinct structures staining with CD200 (Figure S2A) therefore, the majority of CD200 is distinct from the vasculature and lymphatics. Instead, the tubular structures staining for CD200 were found to co-stain for CEA (Carcinoembryonic Antigen) (Figure S2A), which marks secretory and ductal portions of the eccrine sweat gland (3).

In hair follicles, CD200 is expressed by cells with stem-like properties (2). Eccrine sweat glands also contain stem-like cells which contribute to wound healing (4) (5). To determine if CD200 is expressed by the sweat gland stem cells, LGR5 was used as a marker (6) and co-staining for CD200 and LGR5 was observed, demonstrating CD200 expression in sweat gland stem cells (Figure S2A).

Similar to human skin, CD200 is expressed by murine hair follicles. To determine if CD200 is also expressed in murine sweat glands, paw skin, one of the few sites in murine skin to harbour such appendages, was stained for CD200. The CD200 signal was seen in hair follicles in dorsal skin, and in tubular structures in paw skin (Figure S2B). To the best of our knowledge there is no specific murine sweat gland markers available, but this pattern of staining is consistent with eccrine sweat glands at this body site (7). Therefore, CD200 is expressed in eccrine sweat glands, particularly by stem cells.

CD200R1 staining was also tubular in human skin. Therefore, to determine if the same structures express CD200 and CD200R1, co-staining was performed. Figure S2C shows CD200 and CD200R1 co-staining the same structures, with CD200 in ring-like patterns, and CD200R1 staining the ring structure and surrounding cells.

To further decipher the expression of CD200R1, co-staining was performed with markers for lymphatic vessels (LYVE-1), the upper ductal portion of the sweat gland (keratin 10) and the secretory portion of the sweat gland (keratin 18) (8). CD200R1 was observed in lymphatic vessels and the upper ductal portion of the sweat gland, but not in the secretory ducts (Figure S2D). Co-staining with CEA or LGR5 was not possible due to technical difficulties arising from antibodies being raised in the same species.

By flow cytometry, CD200R1 expression is detected in non-haematopoetic cells, and a variety of immune cell types (Figure S1B). To determine if CD200R1-expressing immune cells are in close proximity to sweat glands, immunohistochemistry was performed with CD200R1 and CD45. CD45^+^ immune cells were in close proximity to sweat glands, and a proportion of these immune cells, and those in the epidermis and upper dermis, express CD200R1 (Figure S2E).

**CD200R1 is expressed in naïve and inflamed skin immune cells**

CD200R1 is known to be expressed on immune cells, particularly those of the myeloid lineage. To determine the expression of CD200R1 on skin immune cells, and how this expression changes in psoriasis-like skin inflammation, cells were isolated from ear skin and were analysed by flow cytometry. As expected, CD45^-^ non haematopoetic cells did not express CD200R1. CD200R1 was expressed however, by mast cells (CD45^+^ FcεRIα^+^), macrophages (CD45^+^ F4/80^+^ CD11b^+^), conventional T cells (CD45^+^ CD3^+^ TCRβ^+^) and ILCs (gating as in Figure 4D), but this expression was largely unchanged by psoriasis-like skin inflammation (Figure S3A).

**Differentiation and purity of BMDMs**

BMDMs were differentiated in culture before being stimulated and analysed for activation and cytokine production (Figure 3C-D). To ensure that the cells had differentiated successfully, immediately prior to cell stimulation, samples were taken and analysed by flow cytometry to determine the proportion of macrophages. Cells were typically around 90% F4/80^+^ CD11b^+^ demonstrating adequate differentiation into macrophages had occurred (Figure S3B).

**Table S1: Volunteer data for skin biopsies.**

| **Experimental technique** | **Sample type** | **% Male, % Female** | **Mean age** | **Age range** | **Number** |
| --- | --- | --- | --- | --- | --- |
| **Flow cytometry/IHC** | NN | 43%, 57% | 36 | 21-52 | 7 |
|  | PN | 50%, 50% | 51 | 42-56 | 4 |
| **QPCR** | NN | 11%, 89% | 35 | 21-52 | 9 |
|  | PN | 40%, 60% | 47 | 22-66 | 10 |
|  | PP | 37%, 63% | 45 | 22-66 | 8 |

**Table S2: Antibodies for flow cytometric analysis of human skin.**

|  | **Specificity** | **Conjugate** | **Clone** | **Manufacturer** |
| --- | --- | --- | --- | --- |
| **Panel 1*** | CD200R1 | AF647 | OX108 | Bio-Rad |
|  | CD19 | APCCy7 | SJ25C1 | BD Bioscience |
|  | HLA-DR | eF450 | LN3 | eBioscience |
|  | CD80 | BV510 | L307.4 | BD Bioscience |
|  | CD200 | BV605 | OX104 | BD Bioscience |
|  | CD86 | BV650 | 2331 | BD Bioscience |
|  | CD45 | BV786 | HI30 | BD Bioscience |
|  | CD14 | PECy7 | 61D3 | eBioscience |
|  | CD123 | PECF594 | 7G3 | BD Bioscience |
| **Panel 2** | HLA-DR | FITC | G46-6 | BD Bioscience |
|  | CD45 | PerCPCy5.5 | 2D1 | eBioscience |
|  | CD200R1 | AF647 | OX108 | Bio-Rad |
|  | CD3 | APCCy7 | SK7 | BD Bioscience |
|  | CD66b | V450 | G10F5 | BD Bioscience |
|  | CD127 | BV510 | HIL-7R-M21 | BD Bioscience |
|  | CD200 | BV605 | OX104 | BD Bioscience |
|  | CD117 | BV650 | 104D2 | BD Bioscience |
|  | CD19 | BV711 | SJ25C1 | BD Bioscience |
|  | TCRγδ | PECF594 | B1 | BD Bioscience |
|  | CD56 | PECy7 | CMSSB | eBioscience |

* FITC channel also recorded to measure autofluorescence

**Table S3: Antibodies used for Immunohistochemical staining**

| **Specificity** | **Manufacturer and Clone/cat. No.** | **Secondary reagent used** |
| --- | --- | --- |
| Human CD200R1 | abcam ab198010 | anti rabbit AF546 or AF488 (Invitrogen, A10040 or A11070) |
| Human CD200 | eBioscience OX104 | anti mouse TSA (Invitrogen) |
| Mouse CD200 | eBioscience OX90 | anti rat AF546 (Invitrogen, A11081) |
| Human CD31 | Abcam ab28364 | anti rabbit TSA (Invitrogen) |
| Human CD31 | eBioscience 14-0318-93 | anti mouse AF546 (Invitrogen, A10036) |
| Human LYVE-1 | Abcam ab14917 | anti rabbit AF546 |
| Human CEA | Abcam ab924 | anti rabbit TSA |
| Human LGR5 | Abcam 128242 | anti rabbit AF488 |
| Human CD45 | Abcam ab30470 | anti mouse TSA |
| Human keratin 10 | Abcam ab9026 | anti mouse AF546 |
| Human keratin 18 | Abcam ab668 | anti mouse AF546 |
| Human pDOK1 Y398 | Abcam ab75741 | anti rabbit AF546 |

**Table S4: Antibodies used for flow cytometric analysis of mouse cells**

|  | **Specificity** | **Conjugate** | **Clone/cat. no.** | **Manufacturer** |
| --- | --- | --- | --- | --- |
| **pDOK stain** | pDOK1 Y398 | none | ab75741 | AbCam |
|  | Anti-rabbit | AF488 | A11070 | Invitrogen |
|  | CD45 | BV711 | 30-F11 | BD Bioscience |
| **Neutrophil stain** | CD45 | BV510 | 30F11 | Biolegend |
|  | CD11b | APC or eF450 | M1/70 | Biolegend |
|  | Ly6G  Or Gr1 | PECF594  PECy7 | 1A8  RB6-8C5 | BD Bioscience  eBioscience |
|  | CD200R1 | APC | OX110 | eBioscience |
|  | Annexin V | eF450 | - | eBioscience |
|  | ROS | 520 nm | - | Invitrogen |
| **Macrophages in inflamed skin/BMDM stimulations** | CD45 | BV510 | 30F11 | Biolegend |
|  | F4/80 | BV786 | BM8 | BD Bioscience |
|  | CD11b | APCeF780 | M1/70 | eBioscience |
|  | MHCII | PECy7 | M5/114.15.2 | eBioscience |
|  | CD80 | AF700 | 16-10A1 | eBioscience |
|  | CD86 | eF450 | GL1 | eBioscience |
|  | TNFa | BV650 | MP6-XT22 | BD Bioscience |
|  | IL-12/23p40 | eF660 | C17.8 | eBioscience |
|  | IL-1b | FITC | NJTEN3 | eBioscience |
|  | IL-23p19 | PerCPeF710 | Fc23cpg | eBioscience |
|  | IL-6 | PE | MP5-20F3 | eBioscience |
| **IL-17 production stain** | CD45 | BV510 | 30F11 | Biolegend |
|  | CD11b | FITC | M1/70 | eBioscience |
|  | CD11c | FITC | N418 | eBioscience |
|  | Gr1 | FITC | RB6-8C5 | eBioscience |
|  | F4/80 | FITC | BM8 | eBioscience |
|  | Ter119 | FITC | TER119 | eBioscience |
|  | FcεRIα | FITC | MAR-1 | eBioscience |
|  | CD19 | FITC | 1D3 | eBioscience |
|  | CD3 | PerCPCy5.5 | 145-2C11 | eBioscience |
|  | TCRγδ | PECy7 | GL3 | eBioscience |
|  | TCRβ | AF700 | H57-597 | Biolegend |
|  | Thy1 | BV786 | 53-2.1 | BD Bioscience |
|  | CD127 | BV711 | SB/199 | BD Bioscience |
|  | IL-17 | PEeF610 | eBio17B7 | eBioscience |
|  | CD200R1 | APC | OX110 | eBioscience |

**Figure S1: CD200R1 levels are similar in PN versus NN skin**

NN (healthy) and PN (non-lesional psoriasis) skin punch biopsy cells analysed by flow cytometry. **A.** Cell gating strategy. **B.** Expression of CD200R1 in each population (black line) relative to the isotype control (grey filled histograms). Bar charts n = 4-6. **C.** Fluorescent immunohistochemistry of CD200R1 on NN and PN skin biopsies (n = 4-5). Mean and SD shown. Data were analysed by Mann Whitney test, *p* > .05.

**Figure S2: CD200 is expressed by stem-like cells in the eccrine sweat gland**

Fluorescent immunohistochemistry on abdominoplasty skin to determine the localisation of CD200 and CD200R1. **A.** CD200 was co-stained with markers for the vasculature (CD31), lymphatics (LYVE-1), sweat glands (CEA) and stem cells (LGR5). **B.** Mouse dorsal skin and paw skin stained for CD200. **C.** Co-staining for CD200 and CD200R1 in human skin. **D.** CD200R1 was co-stained with markers for lymphatics (LYVE-1), the upper ductal portion of sweat glands (keratin 10 [K10]) and the secretory portion of sweat glands (keratin 18 [K18]) in human skin. **E.** CD200R1 was co-stained with CD45 to mark immune cells in human skin. Arrow heads indicate cells co-stained with CD200R1 and CD45. Top panel shows sweat glands, bottom panel shows epidermis and upper dermis. Scale bars show 100 μm.

**Figure S3: Flow cytometric analysis of cells**

**A.** Naïve and 3 or 6-day Aldara treated skin cells were analysed for CD200R1 expression by flow cytometry. Grey filled histograms show isotype control staining, black lines show naïve cells, orange lines show cells from skin treated with Aldara for 3 days, red lines show cells from skin treated with Aldara for 6 days. Populations analysed were CD45^-^ non haematopoetic cells, mast cells (CD45^+^ FcεRIα^+^), macrophages (CD45^+^ F4/80^+^ CD11b^+^), conventional T cells (CD45^+^ CD3^+^ TCRβ^+^) and ILCs (CD45^+^ Lineage^-^ CD3^-^ TCRβ^-^ γδTCR^-^ Thy1^+^ CD127^+^). **B.** Representative purity of differentiated BMDMs defined as F4/80^+^ CD11b^+^.

**Supporting Information References**
