## Supplementary figures and images for "Reduced cutaneous CD200:CD200R1 signalling in psoriasis enhances neutrophil recruitment to skin"

### Figure S1

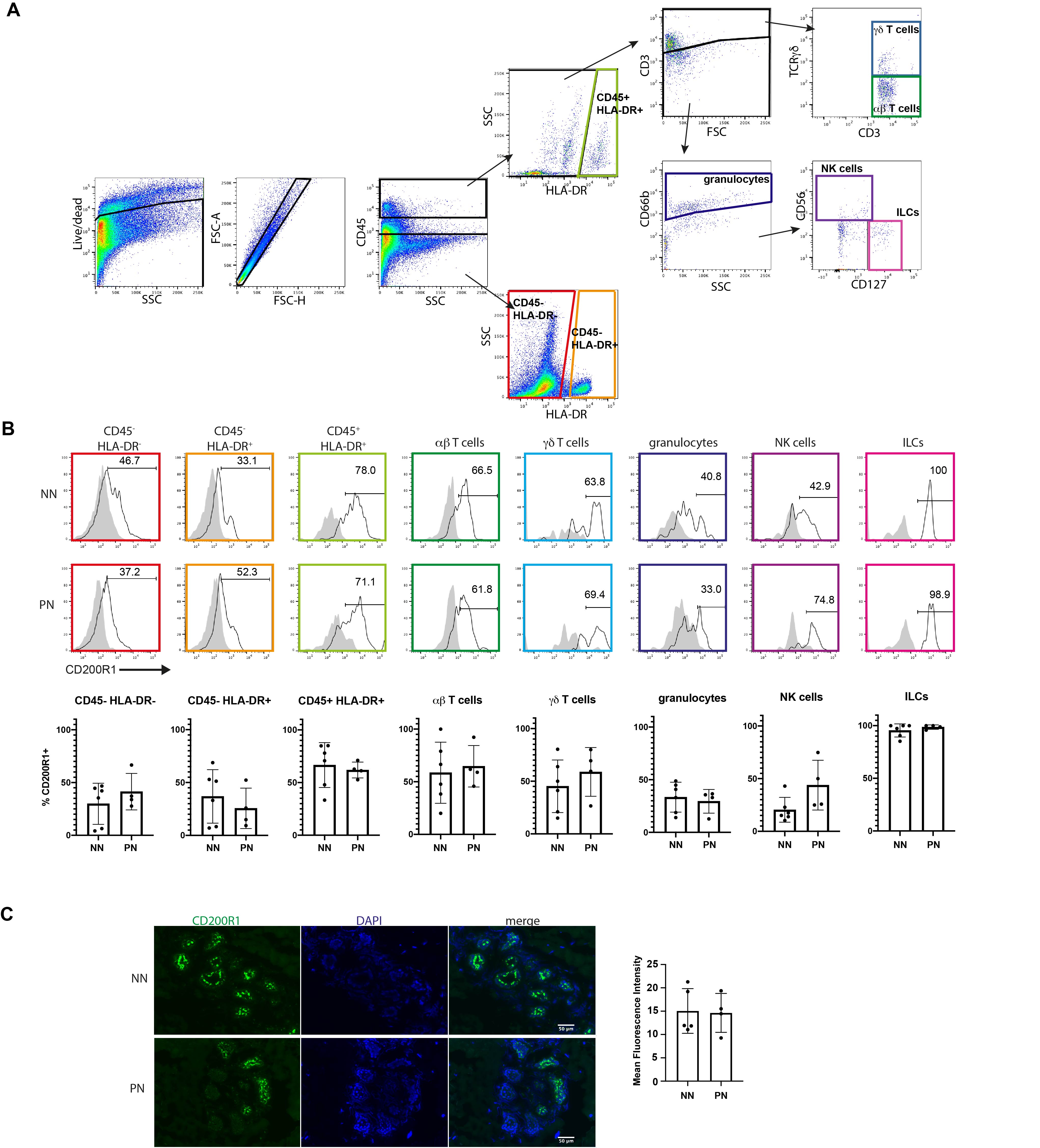

### Figure S2

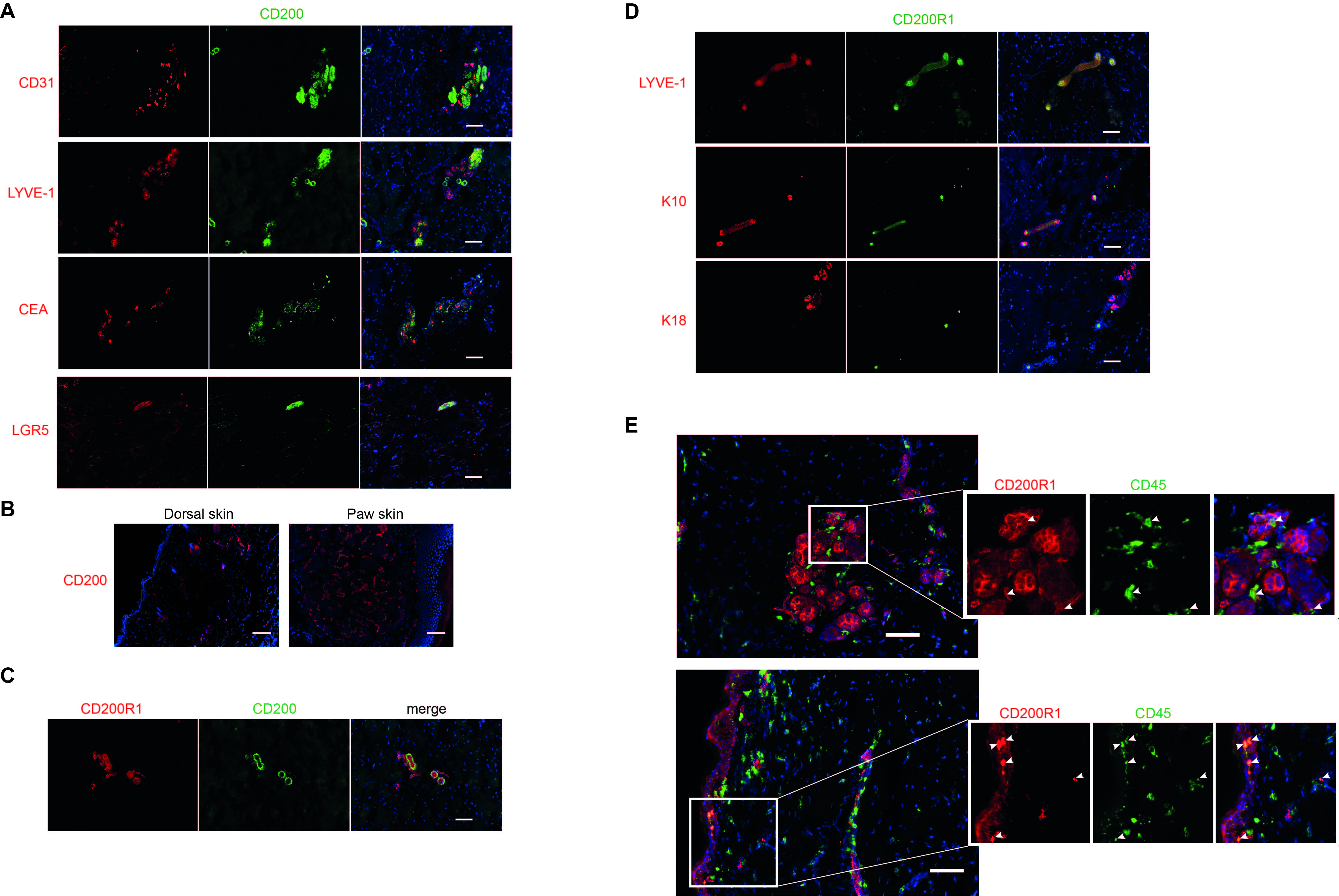

### Figure S3

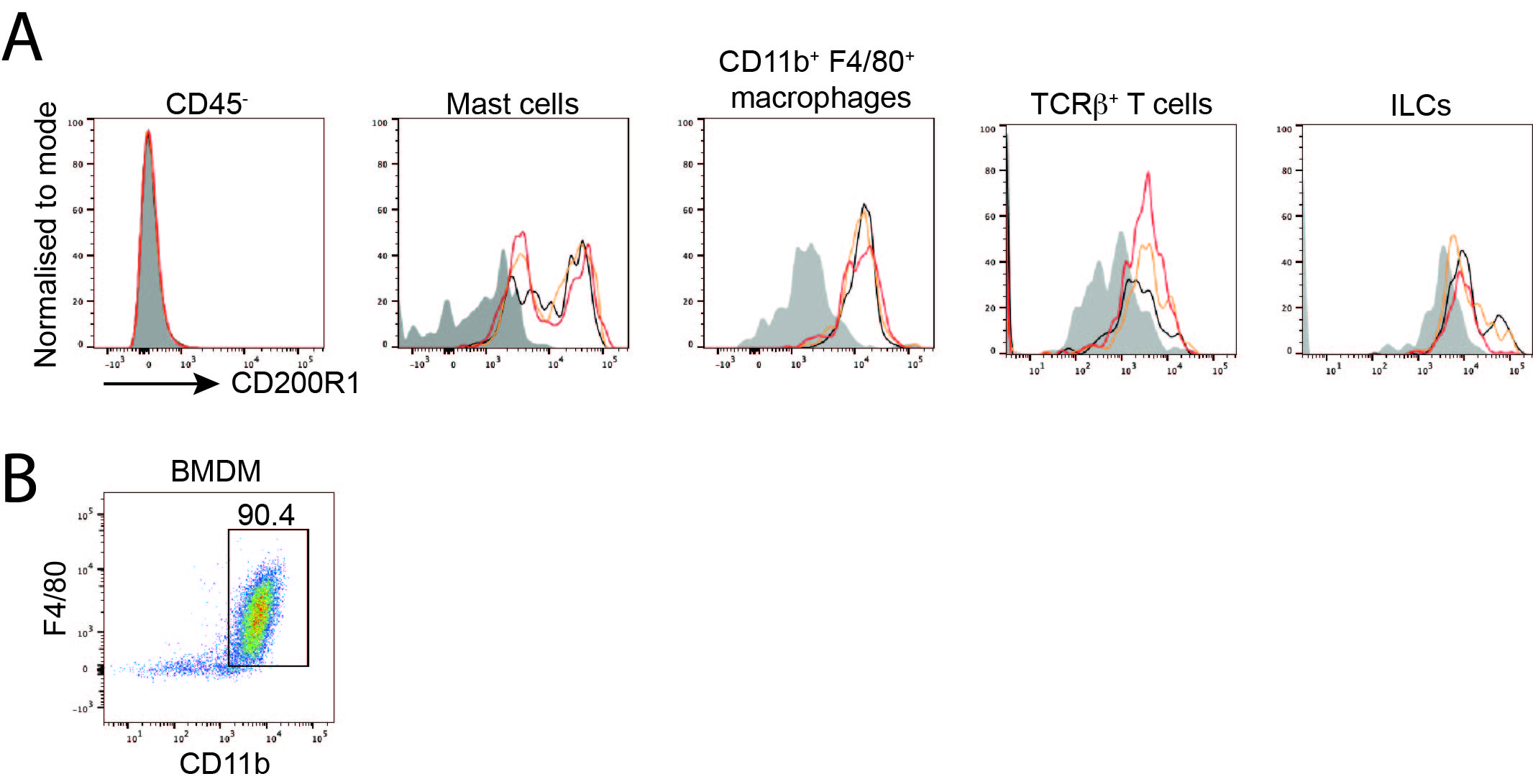
